# Decoding of resting-state using task-based multivariate pattern analysis supports the Incentive-Sensitization Theory in nicotine use disorder

**DOI:** 10.1101/2023.12.31.573585

**Authors:** Cindy S. Lor, David Steyrl, Mengfan Zhang, Feng Zhou, Benjamin Becker, Marcus Herdener, Boris B. Quednow, Amelie Haugg, Frank Scharnowski

## Abstract

**Background:** The Incentive-Sensitization Theory postulates that addiction is primarily driven by the sensitization of the brain’s reward system to addictive substances, such as nicotine. According to this theory, exposure to such substances leads to an increase in ‘wanting’, while ‘liking’ the experience remains relatively unchanged. Although this candidate mechanism has been well substantiated through animal brain research, its translational validity for humans has only been partially demonstrated so far, with evidence from human neuroscience data being very limited.

**Methods:** From fMRI data of N=31 individuals with Nicotine Use Disorder, we created multivoxel patterns capable of capturing wanting and liking-related dimensions from a smoking cue-reactivity task. Using these patterns, we then designed a novel resting-state ‘reading’ method to evaluate how much wanting or liking still persist as a neural trace after watching the cues.

**Results:** We found that the persistence of wanting-related brain patterns at rest increases with longer smoking history but this was not the case for liking-related patterns. Interestingly, such behavior has not been observed for non-temporal measures of smoking intensity.

**Conclusion:** This study provides basic human neuroscience evidence that the dissociation between liking and wanting escalates over time, further substantiating the Incentive-Sensitization Theory, at least for Nicotine Use Disorder. These results suggest that treatment approaches could be personalized to account for the variability in individuals’ neural adaptation to addiction by considering how individuals differ in the extent to which their incentive salience system is sensitized.

## Introduction

The Incentive-Sensitization Theory (IST) has emerged as a prominent framework for investigating the neural processes involved in addiction and dependence to substances. According to this theory, addiction is primarily driven by the sensitization of the brain’s reward system to addictive substances. This leads to an escalating desire or craving for the drug over time, whereas the subjective experience of pleasure or liking remains relatively stable (1,2). The IST has been supported by animal studies showing neuroadaptations in the reward circuitry following repeated exposure to addictive substances (3–6). An essential aspect of the theory is the distinction between psychological dimensions of liking and the wanting. While ‘wanting’ seems to be supported by large dopamine-based regions, ‘liking’ seems more linked to opioids-based systems, and distinct and smaller brain regions, also called hedonic hotspots, such as the ventral pallidum and the parabrachial nucleus (2,7).Although it was mainly built on animal research, the IST has significantly shaped our understanding of addiction in humans. However, the theory’s translation from animal models to human contexts has not been extensively substantiated by empirical evidence, with a limited number of studies corroborating the proposed dissociation between ‘liking’ and ‘wanting’. This dissociation has primarily been demonstrated through behavioral measures such as the Implicit-Association-Test (8,9) and sophisticated behavioral modelling of the transition from a ‘liking’ to a ‘wanting’-dominant incentive (10). Yet, to date, no research using human neuroscientific data has confirmed this dissociation, nor has it been shown that the dissociation intensifies over time.

Recently, functional magnetic resonance imaging (fMRI) has advanced to include multivariate-pattern-analysis (MVPA) methods. MVPA is a machine-learning-based statistical technique that relates spatial patterns of neural activity across multiple brain regions to cognitive or behavioral processes. It can notably be used to decode the neural representations of various stimuli by identifying patterns of activity that are specific to a stimulus of interest or a psychological dimension of interest. For example, MVPA has been employed to decode smoking-related vs. neutral stimuli, with the interesting outcome that the accuracy of the classifier correlated with measures of attentional bias for smoking cues (11). Importantly, MVPA can detect psychological dimensions with greater sensitivity than traditional mass-univariate analyses, which makes this method a suitable candidate tool for dissociating valence-related patterns from craving-related pattern, and which is, in turn, of particular interest for addressing the current gaps in human neuroscientific evidence for the IST.

Our study aims to provide such empirical evidence by leveraging advantages of MVPA which we applied to an existing fMRI dataset (resting-state and cue-exposure task) of cigarette smokers with NUD. Each participant rated stimuli along craving and valence dimensions which serve as analogues to the wanting and liking dimensions, respectively (12). We then developed a multivoxel classification weight-map capable of distinguishing between smoking cues that induce higher craving and those that were rated with higher valence. To complement this analysis, we performed multivoxel regression analyses to identify separate regression weight-maps (13) capable of predicting levels of valence and craving independently. We chose to employ both a classification approach and a regression approach for two main purposes: first, to enhance the overall reliability and robustness of our potential findings. Second, to provide a mitigation strategy against the potential scenario where the instrument measuring craving (or valence) levels surpasses the precision or sensitivity of the instrument measuring valence levels. Such a discrepancy in measurement quality could hinder a direct and meaningful comparison between the two dimensions.

As a second step, we employed a frame-by-frame ‘reading’ method of resting-state scans that immediately followed the task: we applied the classification weight-map and the regression weight-map to each frame of the resting-state scans to estimate how much valence and craving are detectable when the brain is not exposed to any cues.

Finally, we examined the relationship between craving vs. valence brain states at rest and smoking history, as measured by pack-year and years of smoking. In line with the IST, we hypothesize that the pre-post increase of craving pattern expression will escalate with smoking history, while we do not expect to find such correlation for the pattern expression of valence.

## Methods

### Participants and study design

This study is a re-analysis of an fMRI dataset described in (14,15) and consists of N=31 neuroimaging scans of cigarette smokers (age=25.9±5.3; 16F, 14M and 1 non-binary; average daily cigarette consumption=11.5±5.6 cigarettes; Fagerström score=2.8±1.8; years smoking=7.4±4.8; pack-year=4.6±5.2; age of smoking onset=18.5±3.1) who underwent a cue-reactivity task as well as two 7-minute long resting-state scans (one before and one after the task). Inclusion criterium was NUD according to DSM-5. Exclusion criteria were MRI incompatibility, psychiatric or neurological conditions and use of nicotine substitutes.

During the task, the participants passively watched 330 smoking-related pictures of different craving and valence intensities, divided into 5 runs of about 4 minutes each. After scanning, they underwent a picture rating task, during which they rated each stimulus on how much they liked it (valence or ‘liking’ ratings) and on how much it induced craving (craving or ‘wanting’ ratings) (12). Questionnaires on smoking behavior (Questionnaire on Smoking Urges (16) and Fagerström Test for Nicotine Dependence (17) were also administered. Pack-year was defined as daily pack consumption (daily number of cigarettes/20), multiplied by the number of years smoking (18).

### MRI acquisition parameters and preprocessing

MRI acquisition parameters can be found in the supplemental file as well as in (14,15). Resting-state and task data underwent the same preprocessing pipeline using fMRIprep 20.2.6 (19). Resting-state scans were further denoised using independent component analysis (ICA-AROMA, (20). A detailed description of the preprocessing procedure can also be found in the supplemental file.

### Craving vs. valence classification analysis

Our aim is to define a model that classifies high craving pictures vs. high valence pictures. However, we encountered the challenge that some stimuli received similar ratings for both craving and valence, making the distinction between craving and valence difficult. To address this, we implemented this simple selection procedure: we computed the difference between the valence ratings and the craving ratings and sorted them in ascending order. The top 25% of the pictures, exhibiting more valence than craving, were categorized as ‘more valence’, while the bottom 25%, demonstrating a higher craving than valence, were labeled as ‘more craving’. This step discards the pictures with similar craving and valence ratings, and was essential for improving machine-learning performance, since supervised classification analyses are not robust to uncertain or ambiguous labels (21).

We used a support vector machine to classify our brain data. This algorithm was trained using individual activity maps from each subject, a method that has been well-established in the field (22,23). We opted for this method over trial-by-trial brain response analysis to reduce noise levels in our relatively fast-paced design. For each participant, we ran a standard SPM-based first-level general linear model (GLM) analysis (Wellcome Trust Centre for Neuroimaging, London, UK) for which we specified one regressor for all the pictures labelled as ‘more valence’ and one regressor for all the pictures labelled as ‘more craving’. Finally, we defined two contrast maps per subject, one for ‘more valence’ and another one for ‘more craving’. We will refer to them as ‘activation maps’ (Figure 1A). See supplemental file for details.

**Figure 1.**
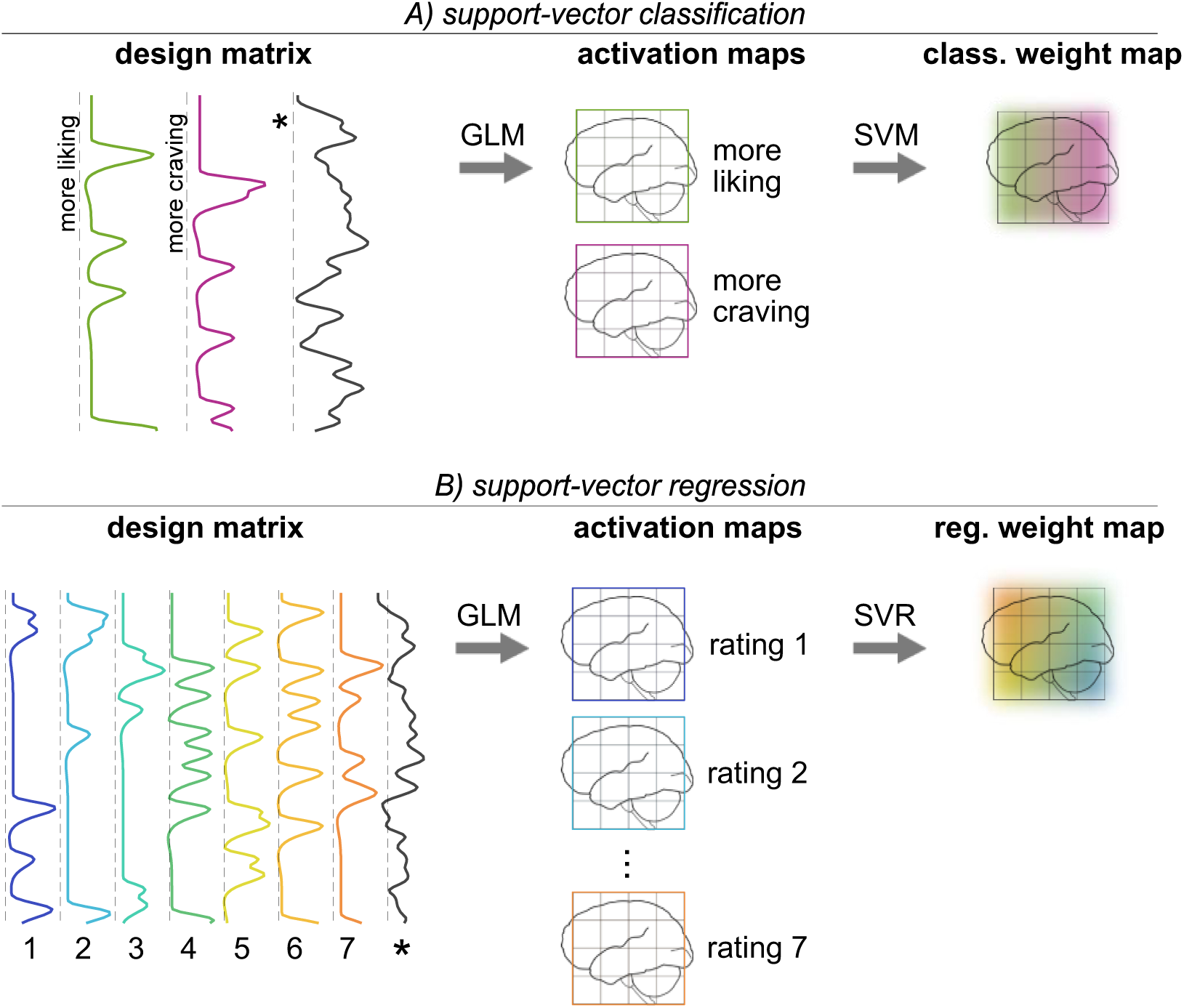
A) Definition of a craving vs. valence classification weight map. Our set of smoking stimuli was divided into two classes: 1) the ones that trigger more valence than craving (valence class) and 2) the ones that trigger more craving than valence (craving class). We ran a GLM analysis with each class being modelled as one regressor to obtain one activation map (or ‘beta map’) per class per subject. B) Definition of independent regression weight maps for craving levels and for valence levels. From the same set of stimuli, we defined one activation map (or ‘beta map’) for seven rating levels per subject and for each dimension separately. The activation maps were then vectorized to serve as input features of a support-vector machine (SVM) (1A) or support-vector regression (SVR) (1B) analysis. *Nuisance regressors

These activation maps then served as the input features of a leave-one-subject-out support vector machine (SVM) analysis. These input features were vectorized and standardized by subtracting the temporal mean value per voxel and dividing each voxel by the total standard deviation. For cross-validation, we partitioned the dataset into N=31 subsets, leaving out one test subject in each subset. We computed an optimal hyperplane from the training set and used the remaining subject as test set. This process was then repeated for each subject in the dataset, allowing us to assess how well the SVM generalizes to predict the target variable for unseen subjects. To evaluate the overall performance of the algorithm, we computed an accuracy measure for each subset and provided the average mean accuracy across all the subsets as a measure of model performance. This step was performed with custom MATLAB scripts (The MathWorks Inc, Natick, Massachusetts, USA) and analysis tools provided by the CanLab (https://canlab.github.io).

Significance levels were assessed using 10000 permutation-based p-values. Each permutation was calculated from shuffling all the labels. See ‘Permutation testing to evaluate model performance’ in supplemental file.

### Valence and craving regression weight-maps

Here, instead of distinguishing valence from craving pictures, our aim is to establish multivoxel patterns capable of capturing valence levels and craving levels separately, i.e., using two independent models. Each picture of the fMRI task was evaluated on both craving and valence dimensions using a fine-grained scale ranging from 1 to 100. To ensure an adequate number of trials per rating level, we opted to discretize the ratings into 7 bins by using a rounding approach (1-7 levels). While binning can result in information loss, this approach allows for a more balanced distribution of trials for each level, which is more robust for the type of analysis we performed.

We ran two separate GLM analyses, one for the craving dimension and another one for the valence dimension, to derive activation maps for each of the seven respective levels. See supplemental file for first-level GLM details. Seven contrast maps per subject and per GLM were then defined, each representing the brain activation associated with each of the seven craving (or valence) levels (Figure 1B).

Next, we used support vector regression (SVR) to obtain a regression weight-map that predicted craving ratings, following a method reported in (23), which used this approach to predict self-reported fear levels. We used a linear kernel (*C* = 1) implemented in the Spider toolbox (http://people.kyb.tuebingen.mpg.de/spider) with normalized individual beta maps (one per rating for each subject) as input features to predict ratings of the grouped pictures. To evaluate the performance of our algorithm, we again used a leave-one-subject-out cross-validation procedure, which means that an optimal hyperplane was computed based on the multivariate pattern of 30 subjects out of 31, and tested on the remaining subject. To evaluate prediction performance, we used overall Pearson correlations (the 7 rating samples of 31 subjects being pooled; 217 pairs in total) and within-subject (7 pairs per subject) between the cross-validated predictions and the actual ratings to indicate the effect sizes, as well as the mean absolute error (MAE) to illustrate overall prediction error. We also ran a permutation test analogous to the classification analysis to evaluate significance levels.

### Pre-vs-Post Resting-state TR-by-TR *‘*reading*’*

We employed a TR-by-TR resting-state ‘reading’ approach to detect the amount of craving vs. valence that can be detected during resting-state, when the brain is not exposed to any smoking cue. Our approach is akin to so-called fMRI ‘replay analyses’ (24,25).

To this end, from the task data, we computed a global classification weight-map from the whole task dataset (Figure 2). From the resting-state data, we excluded the initial five scans of pre and post rest runs to allow for a more stable state, applied a high-pass filter (128s) and removed outlier scans (FD>0.5mm). We concatenated all the functional runs and normalized on a subject-basis by subtracting the mean per voxel and divide by the total standard deviation. We then computed the dot product between the classification weight-map and each frame (each TR) of resting-state scans and added the model intercept (or bias), which resulted in one prediction measure per TR. Next, we averaged these products over all the TRs of the pre-task rest run, as well as for post-task rest run. The resulting values will be referred to as classification pattern expression (PE). Of note, for our classification analysis, a higher PE value indicates more craving than valence and vice-versa.

**Figure 2.**
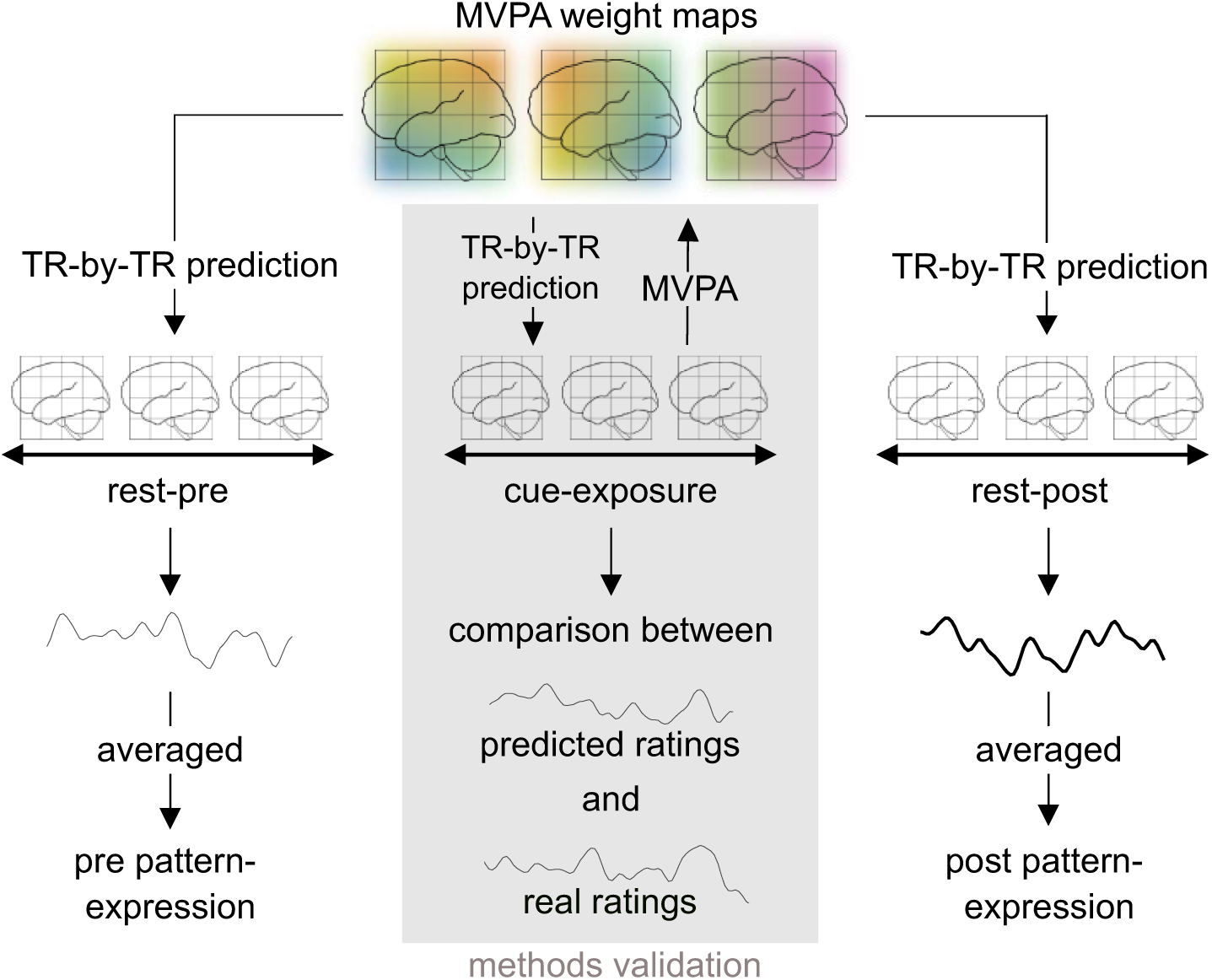
Task-informed ‘reading’ of resting brain states. A smoking cue-reactivity task was used for the definition of multivoxel pattern weight maps that captures craving and valence. The weight maps are then applied to each (pre and post) resting-state frame to evaluate how much this brain pattern is expressed at rest. The same reading procedure was also applied to the task to evaluate how this method performs at predicting single trials.

This reading procedure was repeated using the craving regression weight-map and the valence regression weight-map instead of the classification weight-map (Figure 2). This alternative method enables us to estimate the intensity of craving and valence independently, contrasting with the previous method in which the valence dimension was contingent on the craving dimension. We will refer to the output values as craving-PE and valence-PE respectively. Here, as per intuition, higher PE means higher craving (resp. valence) being detected in the run.

### Trial-Wise Validation of Brain Patterns Derived from Beta Maps

As previously explained, we performed all our MVPA analyses on subject-wise beta maps as a means to train our model on more stable and less noisy brain responses compared to trial-by-trial responses. To ensure that the TR-by-TR reading method still worked at identifying single trials, we ran a trial-wise methods validation analysis. Here, each trial of the task serves as our ground truth (Figure 2, methods validation). For this step, we applied the same ICA denoising procedure to the task scans to ensure comparability with resting-state. To define our true values, we labelled each frame with their corresponding trial type (craving level, valence level or valence vs. craving). As for our predicted values, we computed the dot-product (plus the model intercept) between each frame and the corresponding regression or classification weight-map.

As performance metrics for the regression analyses, we used Pearson correlation between the true values and the predicted values and the mean absolute error. For classification, we used percent accuracy. Since the brain response to a stimulus is expected to occur with the hemodynamic delay, we computed our validation analyses with a temporal shift (from 0 to 9 TRs) between the true and predicted values, with the expectation that the adequation between true and predicted values will progressively peak at a shift of about 6 seconds, which corresponds to a shift between 3-4 TRs. Significance levels were evaluated using permutation testing described in ‘Permutation testing for trial-wise validation’ in the supplemental file.Of note, an additional verification analysis that showed greater task-PE compared to rest-PE (i.e., more stimuli detectability in task) can be found in the supplemental file ‘Task-vs-rest validation analysis’, Figure S1).

### Brain-behavior analysis

We operationalized smoking history using two metrics: pack-years, to estimate cumulative nicotine exposure, and the total number of years spent smoking. This approach is based on the understanding that both the duration and intensity of smoking are important factors, as highlighted by (18). We correlated the pattern expression for craving at rest after the task, corrected for (i.e., by subtraction) the average pattern expression (PE) before the task (i.e., ΔPE), and repeated the procedure for the valence pattern expression. Spearman correlations were used due to violation of normality assumptions. We also correlated the average craving vs. valence classifier output measured at post-task resting-state, corrected for pre-task rest (rest-post minus rest-pre classifier output). We applied Steiger’s Z-test to verify whether the smoking history correlated significantly more with craving-ΔPE than valence-ΔPE.

Again, significance levels were computed using permutation testing by applying the analysis from mock weight-maps so that we obtain a distribution of mock correlations (resp. mock z-values) to which we compare our true correlation (resp. true z-values).

The same analysis was also performed using non-duration smoking variables (Fagerström scores, Smoking Urges scores, daily cigarette consumption and smoking onset) to confirm that our results are mostly driven by time markers and not smoking severity.

## Results

### Validation of classification and regression weight-maps

#### Craving vs. valence classification

On the activation maps, the support vector classifier correctly identified all the maps as either valence or craving (permutation-based chance level=50%; p_perm_<0.0002).

#### Craving regression

The support vector-based regression model predicted craving levels significantly above chance levels, with an overall prediction-outcome correlation of r=0.78 (p_perm<_0.0002); the correlation was done between 217 brain maps (7 levels x 31 subjects) and their respective craving levels. Within-subject prediction-outcome correlation (correlations per subject, then averaged over the 31 subjects) was r=0.88 on average (p_perm<_0.0002). Overall mean absolute error (MAE) was 1.03 (p_perm<_0.0002) and averaged within-subject MAE = 1.03 (p_perm<_0.0002).

#### Valence regression

For valence levels, the support vector-based regression model predicted the ratings significantly above chance, with overall prediction-outcome correlation of r=0.64 (p_perm<_0.0002); the correlation was done between 217 brain maps (7 levels x 31 subjects) and their respective valence levels. Within-subject prediction-outcome correlation (correlation done per subject, then averaged over the 31 subjects) was r=0.77 on average (p_perm<_0.0002). Overall MAE was 1.29 (p_perm<_0.0002) and averaged within-subject MAE = 1.29 (p_perm<_0.0002).

#### Trial-Wise Validation

For all three weight-maps, we achieved the highest prediction performance between true and predicted ratings around the peak of the HRF, which substantiate the validity of our brain reading method (Figure 3). See supplemental file ‘Trial-wise methods validation results’ for details.

**Figure 3.**
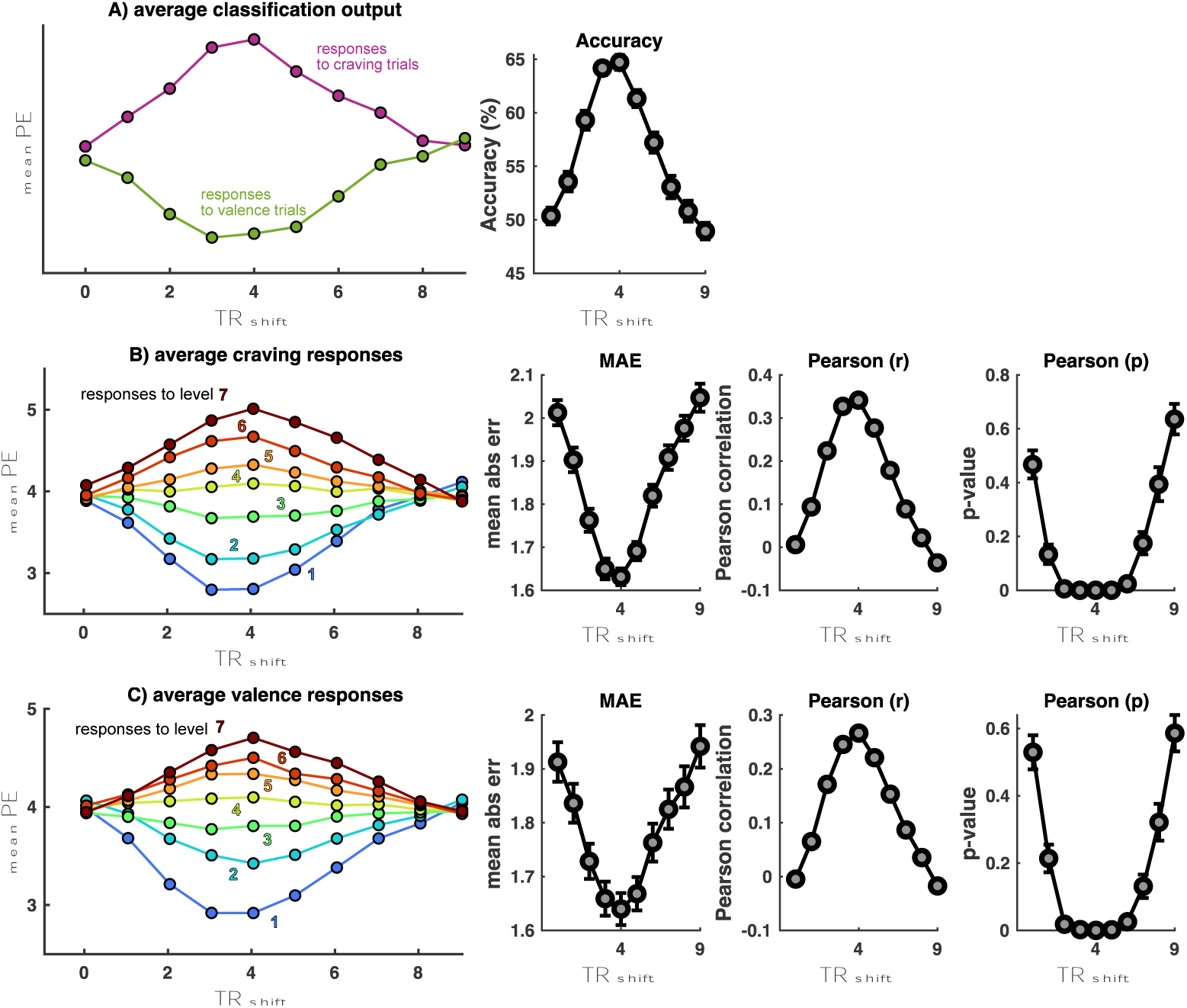
Trial-wise methods validation results. Evaluation of A) the SVM-based craving vs. valence classifier weight map, B) the SVR-based valence regression weight map, and C) the SVR-based craving regression weight map for detecting single-trial levels or class. Hemodynamic response delay was accounted for by including Repetition Time (TR)-based shifts, with maximum prediction performance being expected at the peak of the HRF around 6 seconds (TR=∼3-4). For A, classifier performance was measured with classification accuracy. For B and C, regression performance was measured with mean absolute error (MAE) and Pearson correlation between the predicted rating and true rating.

In summary, our model validation results indicates that all three models can predict ratings or class labels above chance levels. The models were also performant at predicting single trials, and validates them as instruments for TR-by-TR reading of resting-state scans.

### Brain-behavior analysis

#### With Rest Pattern-Expression

As expected, we found that years of smoking correlated with craving-ΔPE (r=0.5707; p_perm_=0.0012) but not with valence-ΔPE (r=-0.1288; p_perm_=0.7427). The Steiger’s Z-test shows that the difference is significant (Z=2.59; p=0.0047). For pack-year, the Steiger’s Z-test also showed that the difference in correlation is significant (Z=1.87; p=0.03), although we did not reach significance for correlations taken separately for craving (r=0.2948; p_perm_=0.0629) and valence (r=-0.1043; p_perm_=0.7160) (Figure 4A).

**Figure 4.**
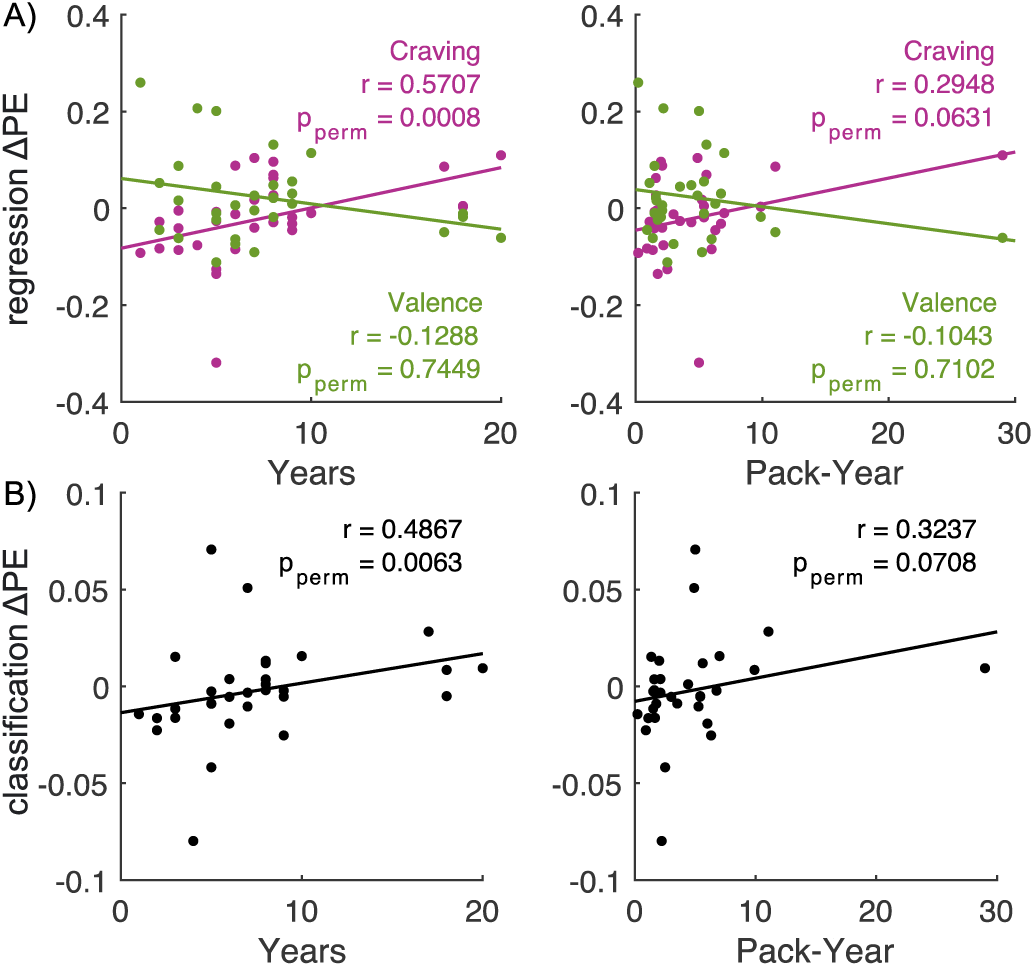
A) Change in craving vs. valence classification output after exposure to smoking cues is linked to smoking history. B) Increase of craving, but not valence, pattern expression (PE) after exposure to smoking cues is linked to smoking history.

Partial correction for age left a significant correlation between craving-ΔPE for Years (r=0.4000; p_perm_=0.0100) but not pack-years (r=0.1342; p_perm_=0.2350). Correlations between valence-ΔPE and Years (r=-0.0254; p_perm_=0.5587) and Pack-year (r=-0.0379; p_perm_=0.5913) remained unsignificant.

Secondly, we correlated classification-ΔPE with both measures with a positive change indicating more craving than valence after the task. We found a significant positive association for Years (r=0.4867; p=0.0055; p_perm_=0.0045) and a positive but not significant one for pack-years (r=0.3237; p_perm_=0.0547) (Figure 4B). Partial correction for age also left a significant correlation between craving-ΔPE for years (r=0.4097; p_perm_=0.0013) but not pack-years (r=0.23; p_perm_=0.0905). The results of the classification are in line with the regression analysis.

Finally, these results did not hold for smoking severity measures that are not time-related (FTND, urge and cigarette consumption). See supplemental file ‘Brain-behavior results with non-temporal smoking markers’ (Figure S2).

## Discussion

The IST suggests that addiction is primarily driven by the sensitization of the brain reward system to addictive substances. A key component is that this sensitization affects how individual experience ‘liking’ and ‘wanting’ of substances: craving increases over time, while the hedonic experience stays stable, or even decreases. This dissociation has been evidenced by animal neuroscientific research (26,27) and human behavioral research (8,28) but the translation to humans has only been partially successful so far (29–31). The present study provides, to our knowledge, the first human neuroscientific evidence that the dissociation between craving and valence escalates over time, from a test sample of NUD individuals. Using advanced neuroimaging techniques and machine-learning approaches on dependent cigarette smokers, our study focused on evaluating the neural persistence at rest of ‘craving’ and ‘valence’ patterns derived from a prior cue-reactivity task.

Here, our patterns were defined using both an SVR-based MVPA procedure (23), to establish regression weight-map capable of predicting craving and valence levels separately, and an SVM-based procedure, to classify ‘higher craving than valence’ from higher valence than craving’ pictures. While SVM operates on a binary classification level, SVR allowed for the identification of patterns associated with continuous levels of craving and valence. The rationale of using both approaches was to first strengthen our findings when results converge, but also to preventively mitigate the hypothetical issue of having one regression weight-map performing significantly better than the other, which might not have allowed for sufficient precision for distinguishing between dimensions.

A strength of our design is the presence of both task and rest within the same session, which allows for higher comparability of within-subject measures, and more importantly, allows to make use of the task time courses as a ground truth to validate our pattern reading method before applying to resting-state. All models achieved higher performance when trained on subject-wise beta-maps than on at the single trial level. At the single-trial level, classification accuracy remained higher than chance and was on par with previous MVPA studies involving similar psychological dimensions (32–34). Regression analyses also achieved high performance for detecting single-trial levels of craving and valence, at similar performance levels as the MVPA prediction of fear levels in (23). Both approaches provided converging conclusions, strengthening the overall validity and reliability of the findings. Our correlation between the neural persistence of craving patterns at rest and the number of years spent smoking, as well as the dissociation between craving and valence trajectories were significant for both measures of smoking history. Interestingly, this finding did not hold for measures of smoking severity that are not time-dependent, which emphasizes the relevance of smoking history on addiction development. Correcting the correlations for age of the participants decreased the value by a significant margin, however, this is not surprising given that the correlation between years of smoking and age was very high (r=0.81). Noteworthy, we did not find significant correlations when we analyzed post-task resting-states alone, which emphasizes the clinical relevance of the psychological changes from pre- to post-task (15).

Conventionally, task-fMRI has been used to decode neural responses evoked by specific tasks or stimuli whereas rest-fMRI is more often used to defined connectivity-based biomarkers associated with psychiatric symptoms or psychological traits. In contrast, our study introduces an innovative combination of brain information obtained from the task to ‘read’, frame-by-frame, resting-state scans acquired during the same session. The interpretation of resting-state from task-derived brain activation has been used in memory research to detect specific replay events of stimuli during resting state (24,36), sometimes even in conjunction with machine-learning-based approaches (35). Here, we expand upon these approaches by applying state-of-the-art task-based classification brain-maps or regression brain-maps to evaluate the underlying psychological content of functional scans even in the absence of explicit cues or tasks, which offers novel perspectives within the field of neuroimaging research in general. Our findings may also inform the customization of therapeutic strategies for substance use disorders. For instance, individuals with longer smoking history who show a pronounced dissociation might benefit more from brain-based interventions specifically designed to address heightened cravings, while short-term smokers could target their hedonic response patterns instead.

As for limitations, it should be noted that, although the dataset employed was suitable for testing our hypotheses, it was not tailored to this particular analysis. To strengthen the study design, one could implement longer interstimulus interval and constraining the ratings scales to a 7-point Likert scale to avoid information loss by rounding. Additionally, the study focused on cigarette smokers, and it remains unclear whether the observed dissociation between craving and valence extends to other substance use disorders. Future research should aim to replicate and expand upon these findings using larger samples and wider distributions of measures of smoking history.

Also, one should note that the instrument for ‘craving’ was slightly better than the instrument for ‘valence’, albeit not significantly, which may have confounded the detection of brain patterns during resting-state. This issue was partly mitigated by performing an additional classification analysis to ensure that our results would also hold using an analogous but distinct analysis. Apart from potential technical or methodological causes, differences in sensor performance may have been psychological. Our stimuli were rated outside the scanner on craving and valence dimensions, and although the cues were likely able to trigger experiences of craving during the rating task, liking, which we called ‘valence’ in this analysis to avoid confusion with the actual hedonic experience, may have involved some form of projection to the presented stimuli, making our liking dimension slightly different from the hedonic experience during smoking itself.

In conclusion, this study takes advantages of advanced machine-learning tools, to provide, to our knowledge, the first human neuroscientific evidence that the dissociation between craving and valence amplifies over time for NUD. Moreover, we introduce an original neuroimaging methodology to decode resting brain states from task-evoked patterns. This dual contribution supplies both the neuroimaging and the addiction research field which are essential for the development of future precision medicine therapeutic tools for substance use disorders.

## Acknowledgments

This research was supported by the Swiss National Science Foundation (BSSG10_155915, 100014_178841, 32003B_166566), the Foundation for Research in Science and the Humanities at the University of Zurich (STWF-17-012), and the Baugarten Stiftung.

## Authors contribution

CSL: Writing-Original draft preparation, Conceptualization, Methodology, Formal analysis. DS: Conceptualization, Methodology, Writing - Review & Editing. MZ: Formal analysis, Data curation. FZ: Software, Methodology, Writing - Review & Editing. BB: Conceptualization, Methodology, Writing - Review & Editing. BBQ: Conceptualization, Writing - Review & Editing. MH: Conceptualization, Writing - Review & Editing. AH: Project administration, Investigation, Data curation, Writing - Review & Editing. FS: Conceptualization, Methodology, Supervision, Funding acquisition, Writing - Review & Editing.

