## Supplemental file for "Decoding of resting-state using task-based multivariate pattern analysis supports the Incentive-Sensitization Theory in nicotine use disorder"

### MRI acquisition parameters

MR scans were collected with a 3T Philips Achieva system (Philips Healthcare, Best, The Netherlands) using a 32 phased-array head coil at the Psychiatric University Hospital, Zurich. Both resting-state and task functional scans were acquired with a T2*-weighted gradient-echo planar imaging (EPI) sequence (repetition time (TR) = 2000ms, echo time (TE) = 35ms, flip angle (FA) = 82°). Resting-state runs consisted of 33 slices, no slice gap, voxel size = 3 x 3 x 3 $mm^3$, field of view (FoV) = 240 x 240 x 99 $mm^3$, total scan duration = 7:12 min per run. Task runs consisted of 27 slices, interslice gap = 1 mm, voxel size = 2 × 2 × 3 $mm^3$ and field of view (FoV) = 220 × 220 × 109 $mm^3$, scan duration = 4 min per run. A high-resolution anatomical T1-weighted scan was also acquired (FA = 8°, 237 slices, voxel size 112 = 0.76 x 0.76 x 0.76 mm3 , FoV = 255 x 255 x 180 $mm^3$ ) at the end of the session.

### MRI Preprocessing

Resting-state scans and task scans were preprocessed using using fMRIprep 20.2.6 (Esteban et al., 2019). Anatomical data preprocessing included field correction, skull stripping, brain tissue segmentation into cerebrospinal fluid, white matter and gray matter, and normalization to MNI space (MNI152NLin6Asym) using nonlinear registration with antsRegistration (ANTs 2.3.3). For each of the five task and two resting-state functional runs, we applied co-registration to the participants' T1-weighted anatomical reference, head-motion parameters estimation, BOLD time-series resampling into native space to correct for head motion (i.e., realignment), then resampling into standard MNI space. For resting-state scans only, automatic removal of motion artifacts using independent component analysis (ICA-AROMA, Pruim et al., 2015) was performed on the preprocessed BOLD on MNI space time-series after removal of non-steady state volumes. Finally, we performed spatial smoothing with an isotropic Gaussian kernel of 6mm FWHM (full-width half-maximum). Corresponding "non-aggressively" denoised runs were produced after such smoothing and several confounding time series (incl. framewise displacement (FD), Power et al., 2012)), were calculated based on the preprocessed BOLD. Task scans were not denoised to avoid removal of signal, but were smoothed with the same kernel in SPM12.

### First-level GLM analysis to generate input feature activation maps

GLMs were performed to generate activation "beta" maps to serve as input features to the machine-learning models. Three separate GLMs were defined in total 1) for craving vs. valence classification, 2) for craving levels regression, and 3) for valence levels regression. For the first one, one regressor of interest was specified per class. For second and third GLM, one regressor was specified for each level (seven regressors of interest per GLM).

Each stimulus was modeled as a boxcar and convolved with the hemodynamic response function to account for the hemodynamic delay. Nuisance regressors of no interest corresponding to the six motion realignment parameters, the first five aCompCor component of the white matter and the first five components of the cerebrospinal fluid, which were computed by fMRIprep, were also added to the model.

Task runs were excluded if maximum framewise displacement (FD) exceeded 5 mm or more than 40% of data points FD > 0.5 mm. Out of the original dataset of 32 participants, one participant was excluded from all analyses due to excess motion in all their runs (N=31), and seven other runs were excluded from five participants.

### Permutation testing to evaluate model performance

To evaluate significance levels, we ran permutation tests as follows: we shuffled the labels within each subject and applied the SVM analysis to predict the shuffled labelled. This permutation analysis was repeated 5000 times, providing a distribution of permuted-based performances, measured in Pearson correlations. Permutation-based p-values were calculated as the number of permuted correlations greater than or equal to the real correlation, divided by the total number of permutations. This provides a quantitative measure of the likelihood that our observed correlation could have occurred by chance alone. We used alpha = 0.05 as significance threshold.

### Permutation testing for trial-wise validation

Permutation testing was also used to assess significance levels of the trial-wise validation analysis. For this, we shuffled all the labels within each subject (ratings for SVR, classes for SVM), computed a 'mock' weight map from all subjects and computed mock performance metrics. We repeated it 10000 times to obtain a distribution of performances to which we compared our true performance (significance threshold = 500/10000 = 0.05). By disrupting the relationship between labels and stimuli, this permutation approach helps to control for potential false positive effects that may arise from unspecific effects not related to the psychological dimension of interest.

### Permutation testing for pre-post behavioral assessment

Permutation testing followed the same principle as the trial-wise validation procedure: 10000 mock weight maps were generated through label shuffling, and each behavioral testing was performed with these mock weight maps to provide a reference distribution to which our correlation values were compared to (significance threshold = 0.05).

### Task-vs-rest validation analysis

As an additional sanity check, verify that more PE can be detected during the actual task (average across all TRs of the five runs) than during resting-state. We tested this using a one-sided paired Wilcoxon signed-rank tests. Again, permutation testing using mock weight maps (10000 iterations) as negative controls were also employed to obtain permutation-based t-values to which we compared our true t-value.

As expected, we detected a significant difference between classification-ΔPE for rest and classification-ΔPE for task which indicates that the task had higher craving content than valence ($p_{perm}$=0.0089). Like for the regression weight map, there was no change in pre-post resting-state PE at the whole-group level.

We were also able to detect higher craving PE during the task (mean PE over the five runs) than resting-state (mean PE over the two runs) ($p_{perm}$=0.0335), but there was no group-level pre-post difference.

Finally, we were also able to detect higher valence PE during the task than resting-state ($p_{perm}$=0.0024), but we found no group-level pre-post difference. See figure S1.

These positive control analyses confirm that we are indeed able to detect more stimuli-related pattern in the task than resting-state.

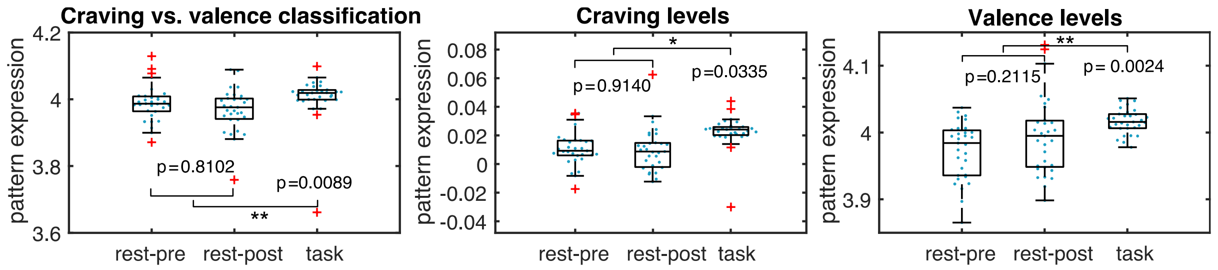

*Figure S1 – Task vs. Rest validation analysis for pattern detection during task compared to rest. Each pattern was applied on a TR-by-TR basis and averaged across rest runs and task runs, to verify whether the tasks showed more pattern expression than resting-state, which was the case (Wilcoxon signed-rank test). There were no pre-post differences at the group level.*

### Trial-wise methods validation results

Craving regression model: the correlation between true values and predicted values is maximum at around the peak of the HRF (4 TRs) ($r=0.34$, $p_{perm}<0.0001$), and the mean absolute error is minimum at the same point ($MAE=1.63$, $p_{perm}<0.0001$) (Figure 3A).

Valence regression model: the correlation between true values and predicted values is maximum at around the peak of the HRF (4 TRs) ($r=0.27$, $p_{perm}<0.0001$), and the mean absolute error is minimum at the same point ($MAE = 1.64$, $p_{perm}<0.0001$) (Figure 3B).

Craving vs. valence classification model: model accuracy peaks at 65% at a +4 TR shift, which corresponds to the peak of the hemodynamic delay. This is significantly better than permutation-defined chance levels (50%; $p_{perm}<0.0001$) (Figure 3C).

These analyses confirm that we are able to detect rating levels (or classes) above chance levels even at the single-trial level and suggest that the weight maps for valence and craving are likely suitable instruments to be used for our resting-state reading method.

### Brain-behavior results with non-temporal smoking markers

We ran an additional analysis to verify that non-temporal smoking variables, Fagerström scores, weekly cigarette consumption, Smoking Urges scores, and age of smoking onset, are less relevant in the context of the Incentive Sensitization Theory. Correlating each variable with craving-ΔPE, valence-ΔPE and classification-ΔPE revealed no significant association (Figure S2).

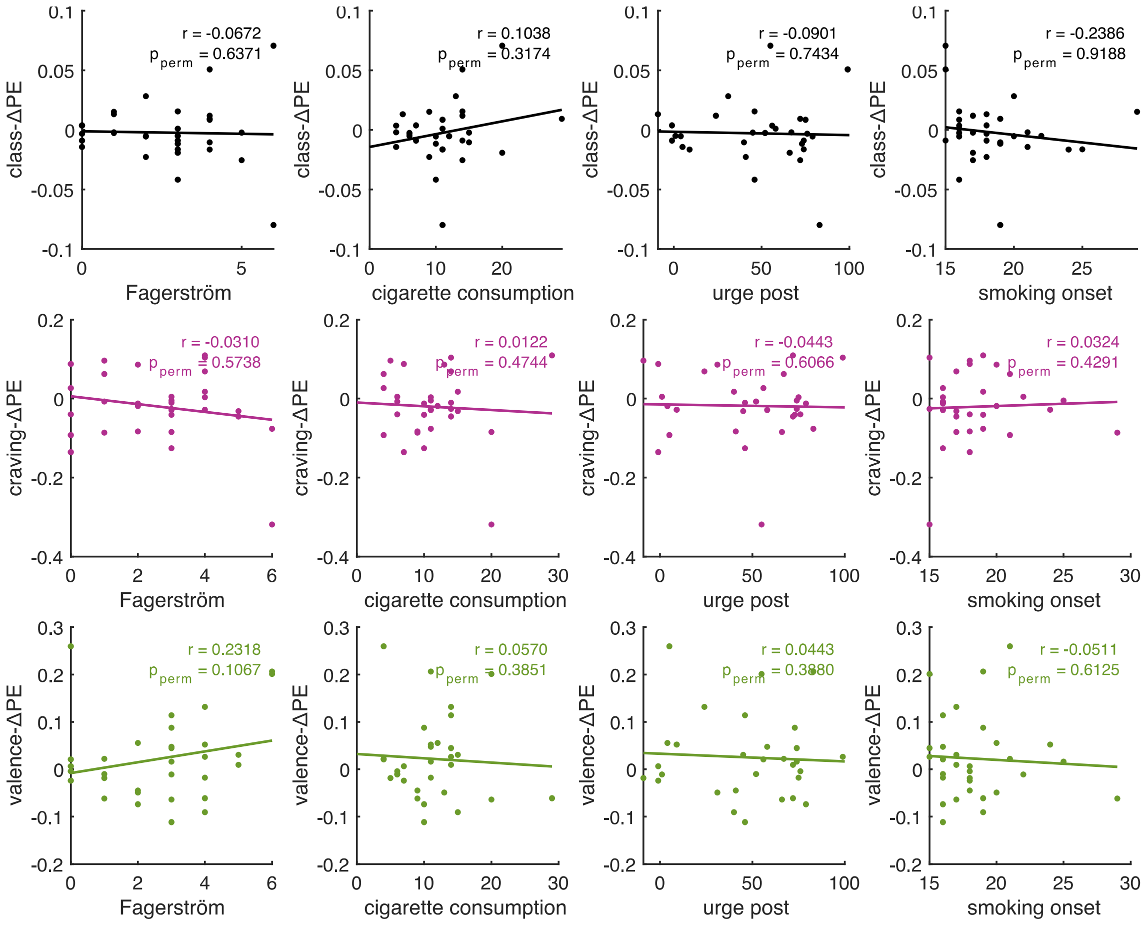


*Figure S2 – Correlations between ΔPE and non-temporal smoking variables.*
